# G-Quadruplex Structures in 16S rRNA Regions Correlate with Thermal Adaptation in Prokaryotes

**DOI:** 10.1101/2024.07.11.603124

**Authors:** Bo Lyu, Qisheng Song

**Author notes:** Corresponding author: Qisheng Song.

## Abstract

G-quadruplex (G4) structure is a nucleic acid secondary structure formed by sequences rich in guanine, playing essential roles in various biological processes such as gene regulation, maintenance of genome stability, and adaptation to environmental stresses. Although prokaryotes growing at high temperatures have higher GC contents, the pattern of G4 structure presence associated with GC content variation in thermal adaptation within genomes and ribosomal genes is rarely reported. In this study, we analyzed 681 bacterial genomes to investigate the role of G4 structures in thermal adaptation. Our findings revealed a strong positive correlation between G4 patterns in the region encoding 16S rRNA genes and optimal growth temperatures (T_opt_), whereas genomic GC content and overall G4 patterns did not show significant correlations with T_opt_. Evolutionary analysis showed significant differences in G4 stability between *Thermotoga* (T_opt_≥ 80 °C) and *Pseudothermotoga* (60°C ≤ T_opt_ < 80°C) species, with *Thermotoga* species exhibiting higher G4 stability, indicating stronger selective pressure for G4 stability under extreme conditions. Circular dichroism analysis showed that specific base mutations at key sites resulted in the absence of G4 thermal stability and structural integrity in *Thermotoga* compared to *Pseudothermotoga*. Collectively, this study suggests that the G4 structures in 16S rRNA encoding regions emerged as key indicators of thermal adaptation and contributes to our understanding of the molecular basis of thermal adaptation.

## Introduction

G-quadruplexes (G4s) are unique nucleic acid secondary structures that form in guanine-rich regions of DNA and RNA ^1,2^. These structures are characterized by the stacking of four guanine bases into a planar arrangement known as a G-tetrad, which is stabilized by Hoogsteen hydrogen bonds ^3^. Multiple G-tetrads can stack on top of each other, forming a stable G4 structure ^4^. The formation and stability of G4 structures are further supported by the presence of monovalent cations such as potassium or sodium, which fit closely between the G-tetrads ^5,6^. G4 structures can adopt various topologies, including parallel, antiparallel, and hybrid forms, depending on the orientation of the DNA or RNA strands and the loop arrangements connecting the G-tetrads ^4,6^. G4 structures play a crucial role in the regulation of gene expression by modulating the transcriptional activity of certain genes ^7,8^. G4 structures are also involved in the maintenance of genome stability, particularly in regions prone to genetic instability, such as telomeres and oncogene promoters ^9–11^. At telomeres, G4 structures protect chromosome ends and regulate telomerase activity, thus playing a vital role in cellular aging and cancer prevention ^10,12^. Additionally, G4 structures are implicated in the replication and transcription processes, where they can act as roadblocks to polymerase enzymes or as binding sites for specific proteins that modulate these processes ^13^. Current research on G4 structures has expanded significantly, revealing their widespread presence and functional importance across various organisms spanning bacteria (e.g., *Escherichia coli* and *Bacillus subtilis*), archaea, and eukaryotes, including humans ^14,15^.

Thermophiles and hyperthermophiles are microorganisms that thrive in extremely high-temperature environments, with thermophiles having optimal growth temperatures (T_opt_) between 45°C and 80°C, and hyperthermophiles exceeding 80°C ^16,17^. These organisms exhibit remarkable structural adaptations that enable their survival and proliferation in such extreme conditions ^18^. Their proteins and enzymes feature increased ionic bonds and hydrophobic cores that prevent denaturation at high temperatures ^19,20^. Membrane lipids of these microorganisms possess unique ether linkages, providing additional thermal stability and preventing membrane fluidity loss ^21^. Additionally, thermophiles and hyperthermophiles have efficient DNA repair systems that mitigate the damaging effects of high temperatures on genetic material ^18,22^. A notable characteristic of these microorganisms is their genomic composition; it was a long debate that the genome of thermophiles and hyperthermophiles displays a high GC content, which contributes to the stability of the DNA double helix ^23,24^. Previous evidence suggests that genomic GC content correlates positively with T_opt_ within prokaryotic families, asserting that environmental factors influencing GC content evolution are less variable among closely related species ^25^. This correlation was observed across 20 prokaryotic families, where the number of families showing positive correlations was significantly higher than expected by chance, irrespective of common ancestry. However, further studies updated T_opt_ values and found that positive correlations between T_opt_ and genomic GC content disappeared in some families ^26,27^. Until recently, this debate appeared to be settled by a large-scale analysis suggesting that prokaryotes thriving in high temperatures exhibit increased genomic GC contents, with thermal adaptation being a proposed reason for this positive association ^28^. However, the results of this study still have limitations, as the positive correlation is weak, and it is evident that some hyperthermophiles, such as *Thermocrinis albus* (46.9%), *Thermodesulfobacterium geofontis* (30.6%), and *Thermotoga petrophila* (46.1%), do not have significantly higher GC contents compared to mesophiles ^28,29^.

Few studies have investigated the relationship between G4 structures and growth temperature in prokaryotic species. One study analyzed the location of putative G4 sequences annotated genomes from the order Thermales, finding these G-rich sequences to be randomly distributed ^30^. Another study speculated that there is no correlation between genomic GC content in Archaea (and Bacteria) and optimal growth temperature, likely because DNA *in vivo* is topologically closed and stable up to at least 107°C ^31,32^. Therefore, a higher density of G4-prone motifs in thermophiles due to a GC-bias is not anticipated ^31^. A comprehensive study of 1,627 bacterial genomes indicated that the highest frequency of G4 forming sequences was detected in the subgroup Deinococcus-Thermus, and the lowest frequency in Thermotogae ^33^. These findings suggest a hypothesis that thermophilic organisms are enriched with G4s as a necessary adaptation to thermally stabilize their genomes for survival at high temperatures, with the underlying evolutionary mechanisms yet to be explored.

In this study, we analyzed 681 prokaryotic genomes from samples covering over a hundred variations of growth temperature to investigate the presence of G4 forming sequences. Our results indicate an evolutionary shift towards an increased frequency and stability of G4s in the region of genome encoding the 16S rRNA genes along the T_opt_ spectrum for the first time. This finding suggests that G4s play a significant role in the adaptation and survival of prokaryotes in high-temperature environments and could be used as one indicator of thermal adaptation for prokaryotes.

## Methods

### Selection and extraction of DNA Sequences

We downloaded the prokaryote growth temperature data (e.g., minimum temperature (T_min_), optimal temperature (T_opt_), and maximum temperature (T_max_)) from the TEMPURA database (http://togodb.org/db/tempura), which contains curated information for 8,645 prokaryotes (549 archaea and 8,096 bacteria). The complete genomic DNA sequences and their corresponding annotation files were obtained from the Genome Database of the National Center for Biotechnology Information (NCBI, https://www.ncbi.nlm.nih.gov/genome). To ensure the reliability and completeness of our dataset, we included only completely assembled genomes in our analysis. Using the links to the NCBI Taxonomy database and the taxonomy IDs provided by TEMPURA for each prokaryotic strain, we selected 681 bacterial genomes for analysis, excluding 155 archaea due to their limited number, which could result in a lack of significant correlations between genomic GC content and T_opt_^28^. The GC contents, genome sizes, and genome accession numbers are provided in Table S1.

Information on 16S rRNA was sourced from the TEMPURA database, and the sequence length and GC content were calculated (Table S2). Prokaryotes were categorized into four groups based on their T_opt_: psychrophiles (T_opt_ < 20 °C), mesophiles (20 °C ≤ T_opt_ < 45 °C), thermophiles (45 °C ≤ T_opt_ < 80 °C), and hyperthermophiles (T_opt_ ≥ 80 °C).

### Identification of G4s in genomic features

QGRS Mapper (https://bioinformatics.ramapo.edu/QGRS/index.php), which uses a predefined motif pattern to identify potential G4 sequences and provides detailed annotations of G4 motifs in specific genes or regions ^34^, was used for the identification of G4 motifs in the 681 16S rRNA encoding sequences. The default parameters for QGRS Mapper were set as follows: QGRS max length: 30, min G-group size: 2, and loop size: from 0 to 36, with a specific loop search string. Genomic G4 distribution patterns were determined by G4Hunter algorithm (https://bioinformatics.ibp.cz), which is more suitable for genome-wide scans ^35,36^. The frequency of G4 motifs was calculated by dividing the number of G4 sequences by the total length. G4 stability was assessed using both the G4Hunter score for genomic sequences and the QGRS score for 16S rRNA sequences.

### Phylogenetic Tree Construction

The exact Taxonomy ID (taxid) for each analyzed group was obtained from the NCBI Taxonomy Database using the Taxonomy Browser. Phylogenetic trees for the 681 bacterial genomes and 12 *Thermotoga/(Pseudo)thermotoga species* ((*Pseudo)thermotoga elfii*, (*Pseudo)thermotoga lettingae*, (*Pseudo)thermotoga profunda*, (*Pseudo)thermotoga hypogea*, (*Pseudo)thermotoga caldifontis*, (*Pseudo)thermotoga thermarum*, *Thermotoga naphthophila*, *Thermotoga petrophila*, *Thermotoga str. RQ2*, *Thermotoga str. RQ7*, *Thermotoga neapolitana*, and *Thermotoga maritima*) were constructed using the Neighbor-Joining (NJ) method. These trees were generated with the “ape” and “phangorn” packages in R (https://www.r-project.org/) by analyzing 16S rRNA gene sequences. The distance matrix was calculated using the Kimura 2-parameter model to ensure accurate representation of evolutionary distances. To assess the reliability and statistical support of the phylogenetic tree branches, bootstrap analysis with one thousand replicates was conducted. The resulting phylogenetic trees, along with their bootstrap support values, were visualized using the Interactive Tree of Life (ITOL) platform (https://itol.embl.de/).

### Relationship between G4s and bacterial growth temperature

We employed phylogenetic generalized least squares (PGLS) regression to examine the relationships among GC contents (whole genome and 16S rRNA), growth temperatures (T_min_, T_opt_, and T_max_), and G4 attributes (frequency and score) using the “caper” package. PGLS enables to detect phylogenetic relationships among species, thus controlling for shared evolutionary history that influences biological traits ^37,38^. Pearson correlation statistics were analyzed to explore the associations between GC content, growth temperatures, and G4 attributes. The *R* value was used to determine the strength of these correlations. The G4 frequency and score across the four groups (psychrophiles, mesophiles, thermophiles, and hyperthermophiles) using ANOVA following normality and Lognormality tests in GraphPad Prism (v5.02, GraphPad Software, Inc.).

### Sequence and LOGO analysis

All 16S rRNA encoding regions for the 12 *Thermotoga/(Pseudo)thermotoga* species were uploaded into UGENE software for comprehensive analysis. The locations of G4 sequences were identified using ClustalW alignment within UGENE (https://ugene.net/). The alignment results were then visualized using Jalview software (https://www.jalview.org/). Each identified G4-forming sequence was analyzed using the RNAfold tool (v 2.6.3) to predict its secondary structure free energy, which was then used as a parameter to assess the stability of the G4s (http://rna.tbi.univie.ac.at//cgi-bin/RNAWebSuite/RNAfold.cgi). To further analyze the sequences, sequence LOGO analysis was generated from the aligned sequences using the WebLogo 3 tool (https://weblogo.threeplusone.com/).

### Circular dichroism (CD) analysis

CD analysis was conducted using a J-1500 CD spectrometer (Jasco International, USA). All spectra were collected within a wavelength range of 220 to 350 nm, with a 1 nm step width and a 1 s response time. The CD spectra represent the average of three scans of the same sample taken at room temperature and are baseline-corrected for buffer signal contributions. DNA oligonucleotide sequences for G4 analysis and their respectively reversed sequences for i-motif analysis were synthesized by Integrated DNA Technologies, Inc. (Coralville, IA, USA) (Table S3). The oligonucleotide sequences (10 µM) used for analyzing G4 structures were heated to 95°C for 10 min in 50 mM Tris buffer (pH 7.5) with or without 100 mM KCl, and then slowly cooled to room temperature over 4 h period. Similarly, DNA oligonucleotide sequences (10 µM) used for analyzing i-motif structures were heated to 95°C for 10 min in 50 mM Tris–acetate buffer at pH 4 and pH 8.1, followed by slow cooling to room temperature over 4 h period.

## Results

### GC Content and Optimal Growth Temperatures in 681 Bacterial Species

We collected a total of 681 bacterial growth temperature datasets using the TEMPURA database and downloaded the corresponding genomic and 16S rRNA gene sequences from NCBI (Table S1 & S2). These 681 bacteria belong to 28 bacterial phyla, including Proteobacteria, Firmicutes, Bacteroidetes, Actinobacteria, Thermotogae, Deferribacteres, Tenericutes, and Deinococcus-Thermus (Fig. 1A). The genome sizes in the dataset range from 800,407 bp to 13,033,779 bp, with genomic GC contents varying from 23.9% to 74.1%. The T_opt_ range from 10°C to 85°C.

**Figure 1.**
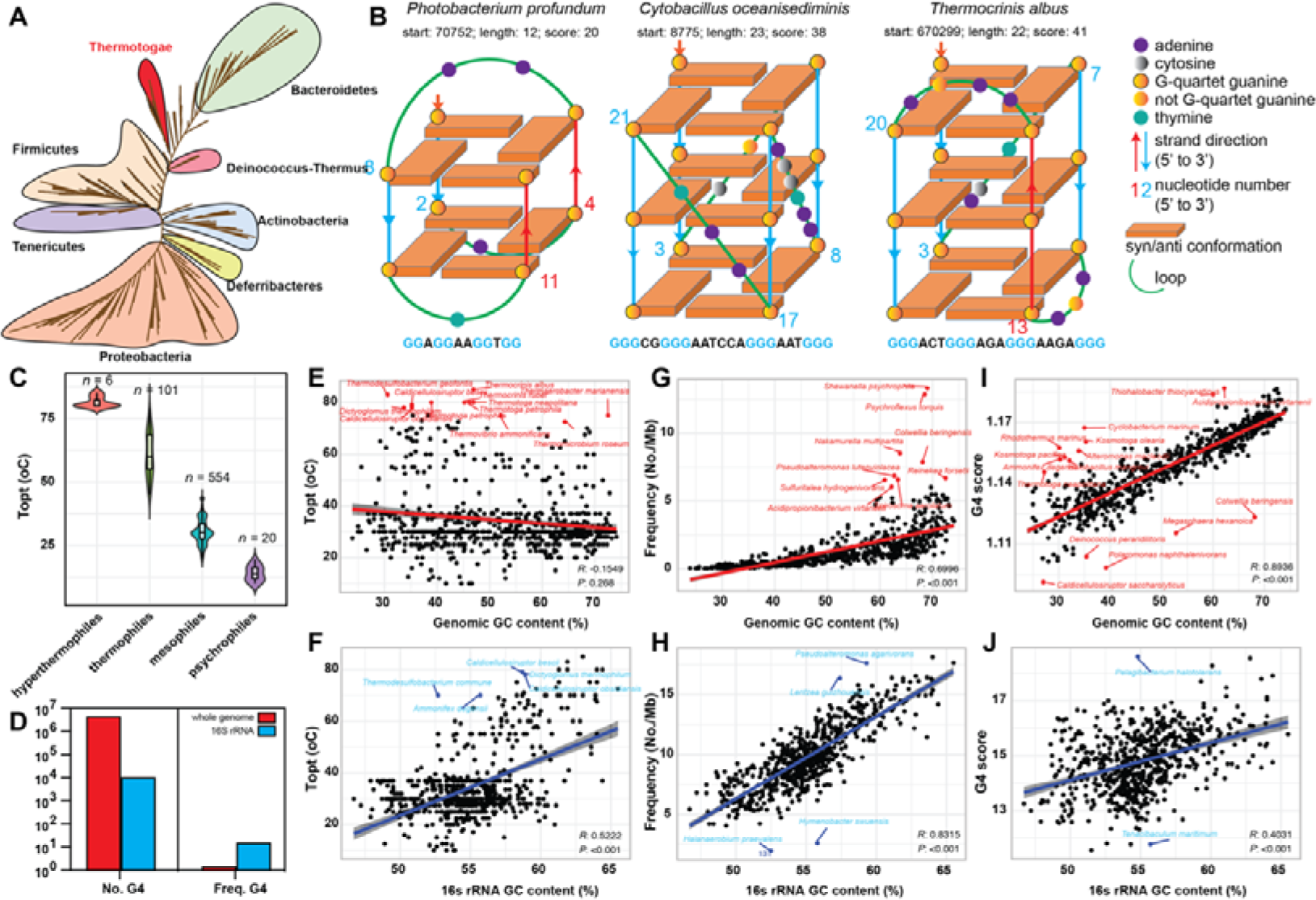
G4 Motif Distribution in 681 Bacterial Species. (A) Unrooted phylogenetic analysis of pathogen genomes based on 681 bacterial species, with different color ranges indicating 28 distinct bacterial phyla. (B) Schematic representation of G4 structures in representative bacterial species, illustrating parallel, anti-parallel, and hybrid arrangements. (C) Classification of bacteria into psychrophiles, mesophiles, thermophiles, and hyperthermophiles based on optimal growth temperatures. (D) Number and frequency of G4 motifs in the genome and 16S rRNA regions. Correlation analysis between bacterial optimal growth temperatures (T_opt_) and the GC content of whole genome (E) and 16S rRNA region (F). Correlation analysis between bacterial GC contents and the frequency and score of G4s in the whole genome (G & H) and 16S rRNA gene region (I-J). PGLS analysis was used to estimate the significance, and Pearson’s *R* value was used to indicate positive or negative correlations.

We used three canonical G4 examples to illustrate the orientation and arrangement of the G-tetrads, along with the loop configurations in the selected bacterial genomes. For instance, anti-parallel, parallel, and hybrid G4 structures are proposed to be present in *Photobacterium profundum* (T_opt_ 10°C), *Cytobacillus oceanisediminis* (T_opt_ 37°C), and *Thermocrinis albus* (T_opt_ 85°C), respectively (Fig. 1B). A high G4 score (e.g., 41 for *T. albus*) can be assigned to structures with three or more G-tetrads, compared to lower G4 scores (e.g., 20 for *P. profundum*) with only two G-tetrads.

### G4 Motif Distribution in Genomes and 16S rRNA Encoding Regions Across Temperature Adaptations

We classified the 681 bacterial datasets into four groups based on their T_opt_: psychrophiles (T_opt_ < 20°C), mesophiles (20°C ≤ T_opt_ < 45°C), thermophiles (45°C ≤ T_opt_ < 80°C), and hyperthermophiles (T_opt_ ≥ 80°C). The majority of bacteria belong to mesophiles (554 species), while the psychrophiles, thermophiles, and hyperthermophiles groups contain 20, 101, and 6 species, respectively (Fig. 1C). We performed comprehensive scans of G4 motifs in both genomes and 16S rRNA encoding regions, identifying a total of 4,160,695 G4 motifs in genomes and 9,794 G4 motifs in 16S rRNA. Interestingly, the frequency of G4 motifs in 16S rRNA (9.79) was significantly higher than in genomes (1.37), indicating a higher potential for G4 formation in 16S rRNA (Fig. 1D). Through PGLS analysis, we identified a very weak positive correlation between genomic GC content and T_opt_, although linear correlation analysis suggested a very weak negative correlation (Fig. 1E). Interestingly, bacteria such as *Thermocrinis*, *Thermotoga*, and *Thermovibrio* have genomic GC contents not exceeding 50%, yet their T_opt_ values are above 70°C. It is unequivocally observed that the GC content of the 16S rRNA was significantly positively correlated with T_opt_ (Fig. 1F). In both genomes and 16S rRNA encoding regions, GC content was positively correlated with G4 frequency and scores, as high GC content facilitates the formation of G4 structures (Fig. 1G-J). PGLS analysis further revealed that only the G4 frequency and scores in 16S rRNA showed a strong positive correlation with T_min_, T_opt_, and T_max_ (Fig. 2A & B), whereas the G4 frequency and scores in genomes exhibited weak or no correlation with these temperature parameters (Fig. S1).

**Figure 2.**
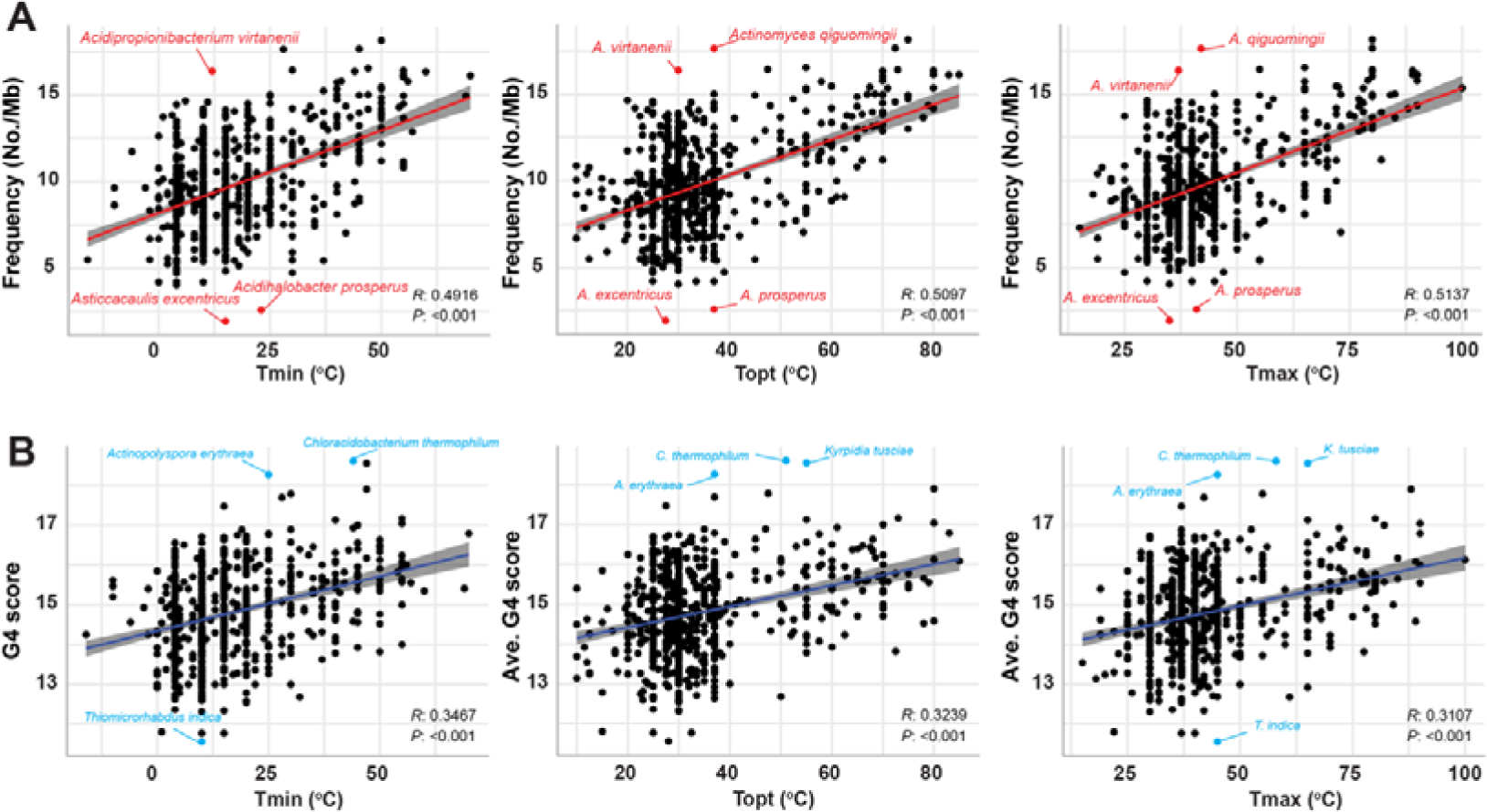
G4 Motif Distribution in Genomes and 16S rRNA Genes. (A) Correlation analysis between bacterial growth temperatures (T_min_, T_opt_, and T_max_) and the frequency of G4s in the 16S rRNA region. (B) Correlation analysis between bacterial growth temperatures (T_min_, T_opt_, and T_max_) and the score of G4s in the 16S rRNA region. PGLS analysis was used to estimate the significance, and Pearson’s *R* value was used to indicate positive or negative correlations.

### Elevated G4 Frequency and Stability in Hyperthermophiles

ANOVA analysis indicated that in the 16S rRNA regions, the G4 frequency in hyperthermophiles was significantly higher than in thermophiles, mesophiles, and psychrophiles (Fig. 3A). Similarly, G4 scores were significantly elevated in hyperthermophiles and thermophiles compared to mesophiles and psychrophiles (Fig. 3B). High-scoring G4 structures are important indicators of G4 stability; we found that 83.3% (5/6) of hyperthermophiles possess three G4-tetrad structures (with scores exceeding 25), compared to 12.9% of thermophiles (13 species) and 2.2% of mesophiles (12 species). Psychrophiles, however, did not exhibit highly stable G4 structures (Fig. 3C & Table S4).

**Figure 3.**
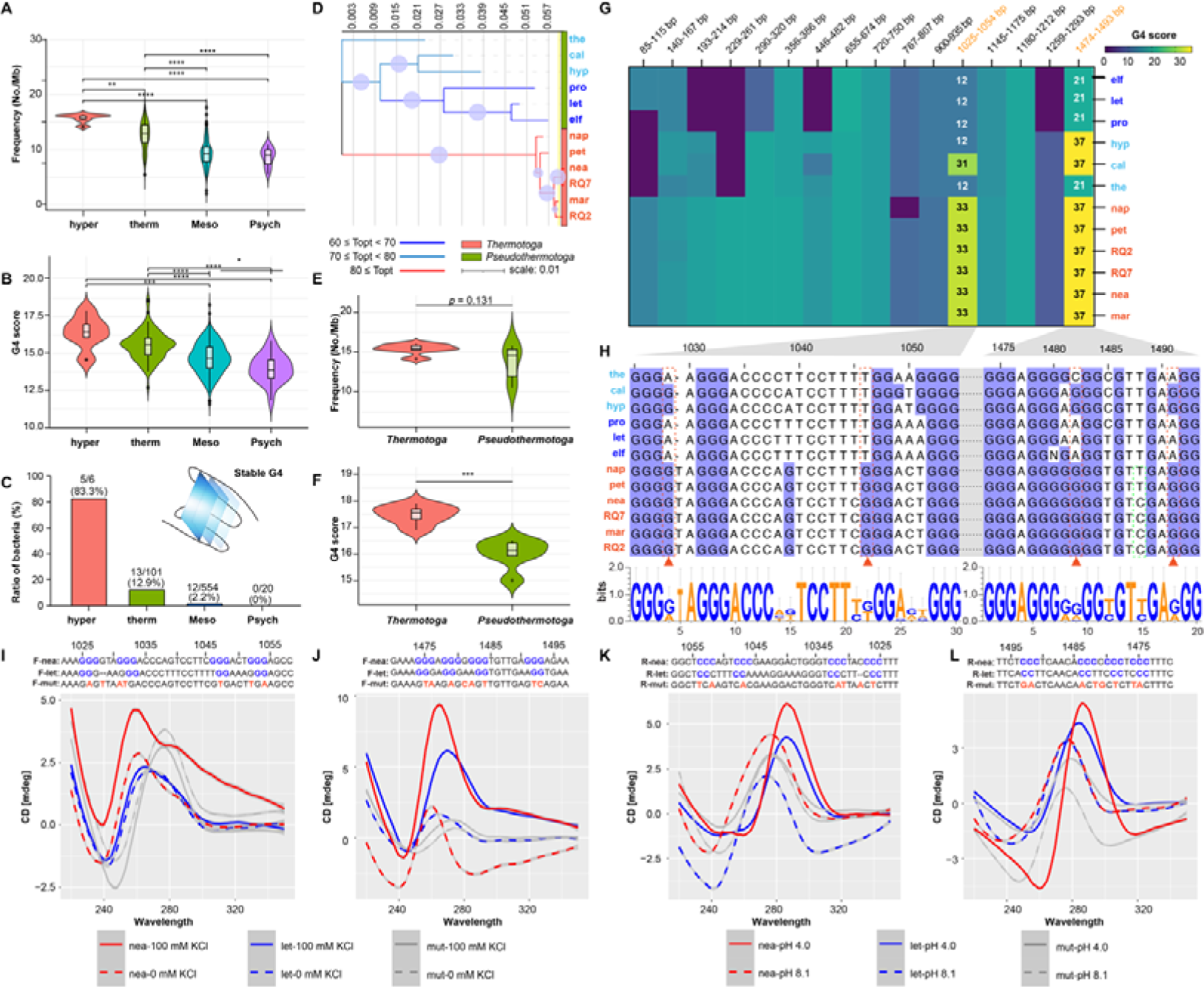
Evolutionary Patterns and Stability of G4 Structures in Hyperthermophiles. (A) Difference in G4 frequency across four groups: psychrophiles (psych), mesophiles (meso), thermophiles (therm), and hyperthermophiles (hyper). (B) Difference in G4 scores across the same four groups. (C) Ratio of stable G4 structures with at least three G-tetrads in psychrophiles, mesophiles, thermophiles, and hyperthermophiles. (D) Evolutionary patterns of 12 species classified into *Thermotoga* and *Pseudothermotoga*. (E) Difference in G4 frequency between *Thermotoga* and *Pseudothermotoga*. (F) Difference in G4 scores between *Thermotoga* and *Pseudothermotoga*. (G) Heatmap showing the presence pattern and scores of 16 G4 structures in *Thermotoga* and *Pseudothermotoga*. (H) Sequence analysis of two stable G4 structures in *Thermotoga*, with LOGO representation of consensus motifs found around positions 1025-1054 nts and 1474-1493 nts in *Thermotoga* and *Pseudothermotoga*. (I-J) Circular dichroism (CD) analysis of the forward ssDNA of the 1025-1054 nts and 1474-1493 nts regions in the absence or presence of 100 mM KCl, showing mutated nucleotides in the mutant sequence. (K-L) CD analysis of the reverse ssDNA of the 1025-1054 nts and 1474-1493 nts regions at pH 4.1 and 8.0, showing mutated nucleotides in the mutant sequence. Abbreviations: nap: *T. naphthophila*, pet: *T. petrophila*, RQ2: *T. str. RQ2*, RQ7: *T. str. RQ7*, nea: *T. neapolitana*, mar: *T. maritima*, elf: *T. elfii*, let: *T. lettingae*, pro: *T. profunda*, hyp: *T. hypogea*, cal: *T. caldifontis*, the: *T. thermarum*, and mut: mutation.

### Evolutionary Patterns and Stability of G4 Structures in the regions encoding 16S RNA

The variability in G4 frequency and scores in hyperthermophiles and thermophiles, prompted further analysis of their potential evolutionary patterns. *Thermotoga* serves as a valuable reference for thermal adaptation analysis due to its well-defined taxonomic classification and the broad range of microorganisms within the genus, exhibiting Topt ranging from 60°C to 85°C. We selected 12 species from the genus *Thermotoga*, all with T_opt_ values above 60°C, based on data from the TEMPURA database and existing literature (Table S5). Phylogenetic analysis supported the current classification of the *Thermotoga* genus, which primarily includes *Thermotoga* (e.g., *T. naphthophila, T. petrophila, T. str. RQ2, T. str. RQ7, T. neapolitana, and T. maritima*) and *Pseudothermotoga* (e.g., *T. elfii, T. lettingae, T. profunda, T. hypogea, T. caldifontis,* and *T. thermarum*) (Fig. 3D). *Thermotoga* species all have T_opt_ values exceeding 80°C, while *Pseudothermotoga* species have T_opt_ values below 80°C. There was no significant difference in G4 frequency between *Thermotoga* and *Pseudothermotoga* (*p* = 0.131) (Fig. 3E), but the G4 scores were significantly higher in *Thermotoga* (Fig. 3F & Table S6). Consequently, we further analyzed the 16 conserved G4 structures expressed in the 12 species to explore specific G4 evolutionary trends and changes. All six *Thermotoga* species possessed these 16 G4s, except for a deletion of one guanine at position 783 nt in *T. naphthophila*, which prevented G4 formation (Fig. 3G & Fig. S2). In contrast, all six *Pseudothermotoga* species had at least two GC rich regions that could not form G4 structures. We found that base mutations are a major factor influencing G4 scores in *Thermotoga* and *Pseudothermotoga*. In at least 11 of the 16 G4 structures, base mutations from adenine, cytosine, or thymine into guanine increased G4 scores in *Thermotoga* (Fig. S2).

Notably, two G4 structures transitioned from unstable forms to stable forms with three G-tetrads, located at 1,025-1,054 nts and 1,474-1,493 nts. Comparing the LOGO sequences at these two positions in *Thermotoga* and *Pseudothermotoga* genomes revealed a conserved G-rich motif across all species (Fig. 3H). However, this “G-richness” is more pronounced in *Thermotoga* species. For instance, guanines at positions 1,028 and 1,046 nts are dominant in *Thermotoga* genomes, while A or T are more common at these positions in *Pseudothermotoga* genomes. Similarly, positions 1,482 and 1,491 nts in *Thermotoga* genes also show near 100% prevalence of guanines. The other 14 G4s showed similar patterns with mutations that have a major/minor effect on G4 formation in *Thermotoga* (Fig. S2). Overall, despite the consensus pattern of favoring G4 formation in both *Thermotoga* and *Pseudothermotoga* species, *Thermotoga* sequences are more conducive to G4 stability, as indicated by higher G4 scores and free energy assessments (Fig. S3).

### CD Spectroscopy Reveals Structural Insights and Stability of G4 and i-motif Structures

We designed three pairs of DNA oligonucleotides for CD analysis: the first pair of sequences comes from *T. neapolitana* representing *Thermotoga* species, the second pair from *T. lettingae* representing *Pseudothermotoga*, and the third pair from mutated sequences (no G4 structure was detected using QGRS Mapper). For the F-strand of the *Thermotoga* group at 1,025-1,054 nts, a maximum peak was observed around 260 nm and a minimum peak at 240 nm (Fig. 3I). When 100 mM K+ was added, the *Thermotoga* group displayed the highest peak, indicating a typical G4 structure ^39,40^. In contrast, the *Pseudothermotoga* and mutated groups showed no change in peak values with or without 100 mM K+, suggesting that neither formed a G4 structure. The *Pseudothermotoga* group’s CD absorption peak was the lowest among all groups due to a base deletion at position 1,028. Mutating the guanine residues at positions - 1,026, −1,028, −1,031, −1,032, −1,047, −1,052, and −1,054 to A, T, A, T, T, T, and A, respectively, caused the maximum and minimum absorption peaks to shift to longer wavelengths, suppressing the K+-mediated enhancement of peak intensity compared to the *Thermotoga* group (Fig. 3I). Similarly, for the G4 structure at 1,474-1,493 nts, the addition of 100 mM K+ enhanced the absorption peaks in both the *Thermotoga* and *Pseudothermotoga* groups (Fig. 3J). However, in the mutant (with guanine residues at positions −1,475, −1,476, −1,479, −1,481, −1,482, −1,484, −1,493, and −1,494 mutated to T, A, A, C, A, T, T, and C, respectively) and under 0 mM K+ conditions, no enhancement of absorption peaks was observed (Fig. 3J).

For the R-strand of the *Thermotoga* group at 1,025-1,054 nts, the highest and lowest absorption peaks were around 290 nm and 260 nm, respectively (pH 4.0), with the *Thermotoga* group exhibiting a significantly higher peak than the *Pseudothermotoga* group (Fig. 3K). At pH 8.1, the maximum and minimum peaks shifted to 280 nm and 250 nm, respectively, consistent with previous analyses of i-motif structures by CD spectroscopy ^40,41^. Corresponding mutations also caused the maximum and minimum absorption peaks to shift to shorter wavelengths, blocking the H+-induced enhancement of peak intensity. Similarly, for the i-motif structure at 1,474-1,493 nts, acidic conditions induced the formation of i-motif structures in both the *Thermotoga* and *Pseudothermotoga* groups, with the *Thermotoga* group showing significantly higher absorption peaks than the *Pseudothermotoga* group (Fig. 3L). Non-acidic conditions (pH 8.1) and mutant did not form i-motif structures.

## Discussion

Our study provides a comprehensive analysis of the relationship between G4 structures and growth temperatures in prokaryotes by examining 681 bacterial genomes. This endeavor was inspired by the longstanding debate about the role of genomic GC content and its impact on thermal stability and adaptation. Our PGLS analysis suggested a very weak correlation between GC content and T_opt_, highlighting the complexity of thermal adaptation and suggesting that while GC content might play a role, it is not the sole factor. For example, species like *Thermocrinis albus*, *Thermotoga neapolitana*, and *Thermovibrio ammonificans* thrive at temperatures above 70°C despite having GC contents below 50%, indicating that other genomic features or regulatory mechanisms might significantly contribute to thermal stability. Therefore, despite the recent study claiming to resolve the previous contradictory observations and end the long debate by stating that prokaryotes growing at high temperatures have higher GC contents ^25–28,42,43^, but this correlation is very weak and likely applies only within certain temperature ranges, such as those of psychrophiles and mesophiles but not thermophiles and hyperthermophiles. Additionally, the relationship between GC content and growth temperature may be influenced by other significant evolutionary forces ^44,45^. For example, in the genus *Thermotoga* (Topt range 60-85°C), the GC content ranges from 38.5% to 51.5%. Their relatively low GC content could be due to genome reduction and the loss of DNA repair genes, leading to decreased DNA repair efficiency ^46,47^. Despite this, the frequency of G4 structures in *Thermotoga* is high, likely playing a significant role in maintaining genome integrity and contributing to thermal adaptation.

Furthermore, our research indicates that while the G4 patterns in 16S rRNA encoding regions are strongly associated with T_opt_, the genomic G4 patterns do not show a significant correlation with T_opt_. This distinction is crucial as it points to a more specialized role of G4 structures in 16S rRNA in thermal adaptation, which is particularly relevant for hyperthermophiles and thermophile, where the need for stable ribosomal function is paramount due to the extreme conditions they inhabit ^48,49^. The 16S rRNA is essential for ribosome assembly and function, processes critical for protein synthesis and, consequently, for cell survival ^50^. Formation of stable RNA G4 has been reported in large ribosomal RNA, where the authors suggested that these G4s are a common feature of the large subunit rRNA in human genome, potentially serving as switches between inter- and intramolecular G4s in rRNA tentacles ^51^. Comparative studies have highlighted the role of G4 structures in the regulatory regions of genes in thermophilic archaea and bacteria, supporting the idea that these structures help regulate gene expression under thermal stress ^31,33^. Our findings add to this body of knowledge by specifically identifying the role of 16S rRNA G4 structures in thermal adaptation. This specificity is crucial because it points to a targeted adaptation mechanism where G4 structures in 16S rRNA enhance the stability and functionality of ribosomes at higher temperatures. The presence of stable G4 structures in 16S rRNA likely provides a selective advantage to bacteria in extreme environments ^52^. These structures might help maintain the integrity and efficiency of ribosome function under thermal stress, ensuring that protein synthesis can proceed uninterrupted ^53,54^. This is particularly important for hyperthermophiles, which thrive at very high temperatures and need highly stable molecular structures to survive. The elevated G4 frequency and stability in these organisms suggest a strong selective pressure favoring the formation of G4 structures in key genomic regions like those encoding the 16S rRNA genes.

Nature acts as a masterful and prolific chemist, creating numerous hostile niches that serve as ideal habitats for various thermophiles ^17^. Among these, subsurface environments are particularly significant for the primary domain of the bacterial genus *Thermotoga* ^55^. To date, more than 60 species of thermophilic bacteria and archaea have been identified, with certain species of the genus *Thermotoga* being of utmost importance ^56^. Phylogenetic analysis supported the current classification of these genera, which was proposed to be split into two genera *Thermotoga* and *Pseudothermotoga* ^57^, and this classification was further corroborated by differences in G4 structure stability between the two groups. While there was no significant difference in the frequency of G4 structures between *Thermotoga* and *Pseudothermotoga*, the significantly higher G4 scores in *Thermotoga* suggest a stronger selective pressure for G4 stability in these high-temperature environments. Bacteria that thrive in high-temperature environments, such as those in the Deinococcus-Thermus group, are highly abundant in G4 structures ^30^. This abundance is also related to their adaptation to other stress factors, such as radiation ^58^. These stable G4 structures could help prevent DNA damage, facilitate efficient DNA replication and transcription, and ensure proper cellular function despite the high temperatures ^13^. In contrast, the *Pseudothermotoga* or even mesophile species, which thrive in relatively cooler environments, may experience limited selective pressure for G4 stability, leading to differences in the structural properties and functions of their G4 motifs. For example, in the mesophilic bacterium *E. coli*, which has an optimal growth temperature of around 37°C, G4 structures are less prevalent and often less stable compared to those in thermophiles ^15,59^.

Our detailed analysis of 16 conserved G4 structures across these species revealed specific mutations that impact G4 formation. *Thermotoga* species consistently expressed these G4 structures, while *Pseudothermotoga* species showed multiple non-forming G4 structures due to specific mutations. Specifically, the G4 structures in *Thermotoga* species maintained intact G-tetrads at several key positions, such as guanine residues at positions 1,028 and 1,046. In contrast, these positions in *Pseudothermotoga* species were often replaced by other bases, such as adenine or thymine, disrupting the formation of G4 structures. For example, mutations in *Pseudothermotoga* at critical guanine positions to other nucleotides led to the loss of G4 stability, preventing the formation of stable G4 structures. This evolutionary divergence indicates that *Thermotoga* species may have developed stable G4 structures as a response to high-temperature environments. Similar to the Levy jump model ^60^, there is an accidental jump in the evolution rate of GC content and growth temperature. The jump of GC content is significantly correlated with the change of growth temperature, and the formation of G4 structure may be induced during the skip process ^28,60^. On the other hand, *Pseudothermotoga* species, living in relatively cooler environments, may not experience the same selective pressure to maintain G4 stability.

The conservation of G4-forming sequences and their functional relevance across species implies selective pressures that preserve these structural motifs throughout evolution. Similarly, the ancient sequencing of Hepatitis B virus (HBV) revealed its long-standing association with humans, and analysis of G4 sequences in both ancient and modern HBV genomes showed a convergence in G4 frequencies with their hosts, suggesting an evolutionary “genetic camouflage” strategy aiding in viral persistence and evasion of host defenses ^61^. Furthermore, studies have revealed variations in G4 structures among species, including differences in sequence motifs, loop lengths, and structural stability, reflecting adaptations to specific genomic contexts and cellular environments ^62,63^. For example, over 5 million gains/losses or structural conversions of G4s can be caused by single-nucleotide variations in human genome, affecting transcription factor-binding sites and enhancers ^62^. The evolutionary dynamics of G4s also extend to their interactions with regulatory proteins, nucleic acids, and small molecules, shaping complex regulatory networks and signaling pathways ^14,64^.

Our study underscores the critical role of G4 structures in the thermal adaptation of bacteria, particularly through their presence and stability in 16S rRNA genes. The strong correlation between G4 stability and high growth temperatures suggests that these structures play a vital role in enabling bacteria to survive and thrive in high-temperature environments. This finding has significant implications for our understanding of bacterial adaptation and evolution. Future research should focus on experimentally validating the functional roles of G4 structures in vivo. This could involve exploring the impact of G4 structures on gene regulation, genome stability, and overall cellular function under different environmental conditions. Additionally, investigating the interplay between G4 structures and other genomic features, such as DNA repair mechanisms and regulatory proteins, could provide deeper insights into the molecular basis of thermal adaptation.

### Conclusion

In conclusion, our research underscores the pivotal role of G4 structures in the thermal adaptation of prokaryotes, with a particular emphasis on the 16S rRNA genes in thermophilic species. The strong positive correlation between G4 patterns in these genes and T_opt_ highlights their importance in maintaining ribosomal stability and function under extreme thermal conditions. The evolutionary analysis of *Thermotoga* and *Pseudothermotoga* species revealed significant differences in G4 stability, suggesting that stable G4 structures provide a distinct adaptive advantage in high-temperature environments. Other than 16S rRNA GC content, we also consider G4 patterns as one of the key indicators for the growth temperature of prokaryotes. Overall, our findings contribute to a deeper understanding of the molecular mechanisms underpinning thermal adaptation and offer promising directions for future research and innovation.

## Supporting information

Graphic abstract

## Acknowledgment

None.

## Credit author statement

Bo Lyu: writing original draft, data analysis, and visualization. Qisheng Song: writing original draft, reviewing and editing.

## Ethical approval

Ethical review and approval were not required for this study.

## Availability of data and materials

The original temperature datasets analyzed in this study are available from the TEMPURA Database on prokaryotic growth temperatures (http://togodb.org/db/tempura). Supplementary materials accompanying this article provide further data and detailed information that support the study’s findings and conclusions. Researchers and interested individuals can access these materials to explore and validate the study results.

## Consent for publication

Not applicable.

