## Supplemental figure s1-s3 for "G-Quadruplex Structures in 16S rRNA Regions Correlate with Thermal Adaptation in Prokaryotes"

Bo Lyu and Qisheng Song*

Division of Plant Science and Technology, University of Missouri, Columbia, MO 65211, USA

*Corresponding author:

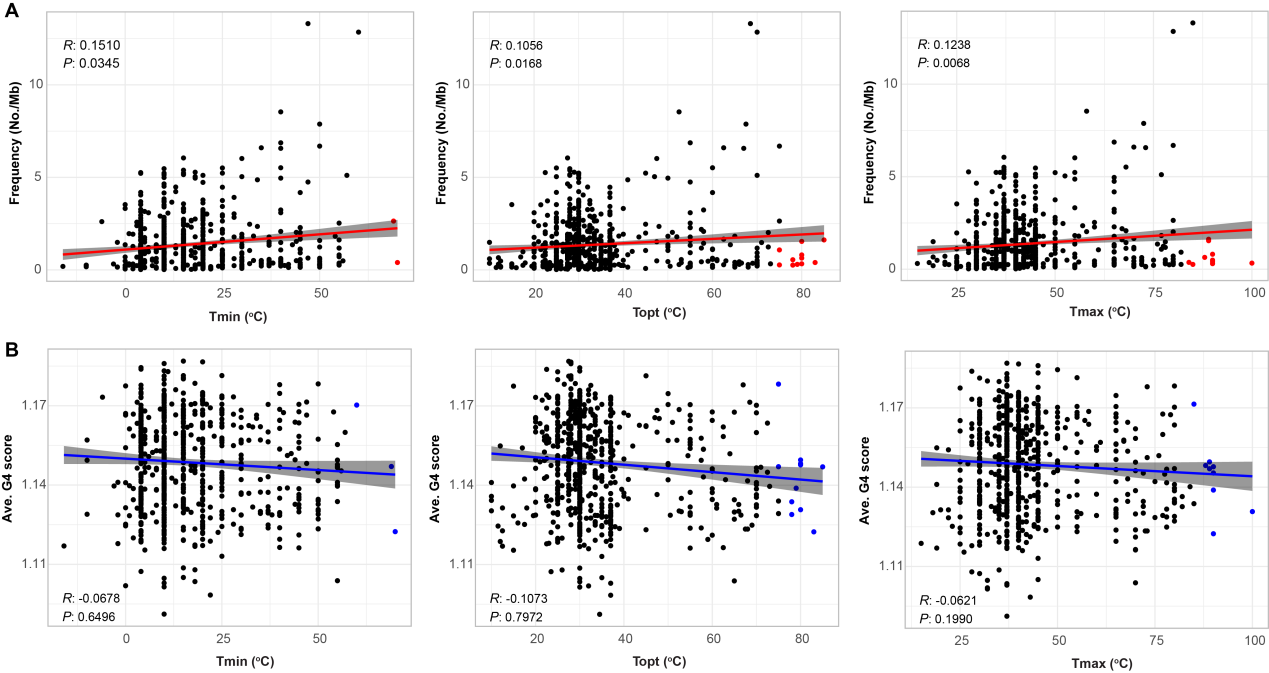

**Figure s1. G4 Motif Patterns in Genomes and Their relationship with Growth Temperatures.** (A) Correlation analysis between bacterial growth temperatures (T_min_, T_opt_, and T_max_) and the frequency of G4s in the whole genome. (B) Correlation analysis between bacterial growth temperatures (T_min_, T_opt_, and T_max_) and the score of G4s in the whole genome. PGLS analysis was used to estimate the significance, and Pearson’s *R* value was used to indicate positive or negative correlations.

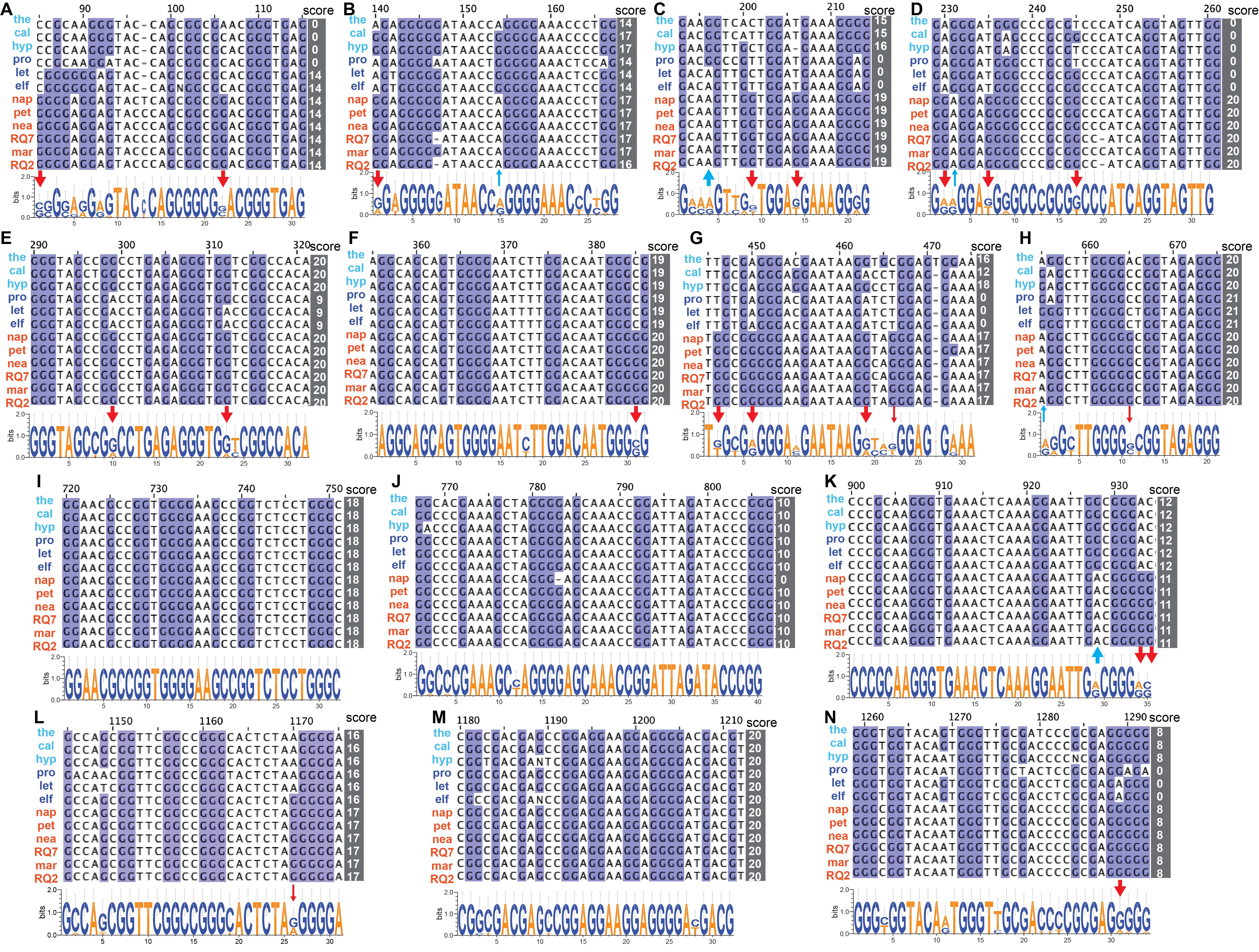

**Figure s2. Sequence analysis and LOGO visualization of 14 G4 structures in *Thermotoga* and *Pseudothermotoga*.** Sequence analysis and LOGO visualization of 14 G4 structures in Thermotoga and Pseudothermotoga. The 14 G4 consensus motifs are located at positions 85-115, 140-167, 193-214, 229-261, 290-320, 356-386, 446-482, 655-674, 720-750, 767-807, 900-935, 1145-1175, 1180-1212, and 1259-1293 nts. Red bold/regular dashes indicate guanine mutations (from adenine, cytosine, or thymine) that have a major/minor effect on G4 formation in *Thermotoga*. Blue bold/regular dashes indicate guanine mutations that have a major/minor effect on G4 formation in *Pseudothermotoga*. Abbreviations: nap: *T. naphthophila*, pet: *T. petrophila*, RQ2: *T. str. RQ2*, RQ7: *T. str. RQ7*, nea: *T. neapolitana*, mar: *T. maritima*, elf: *T. elfii*, let: *T. lettingae*, pro: *T. profunda*, hyp: *T. hypogea*, cal: *T. caldifontis*, the: *T. thermarum*, and mut: mutation.

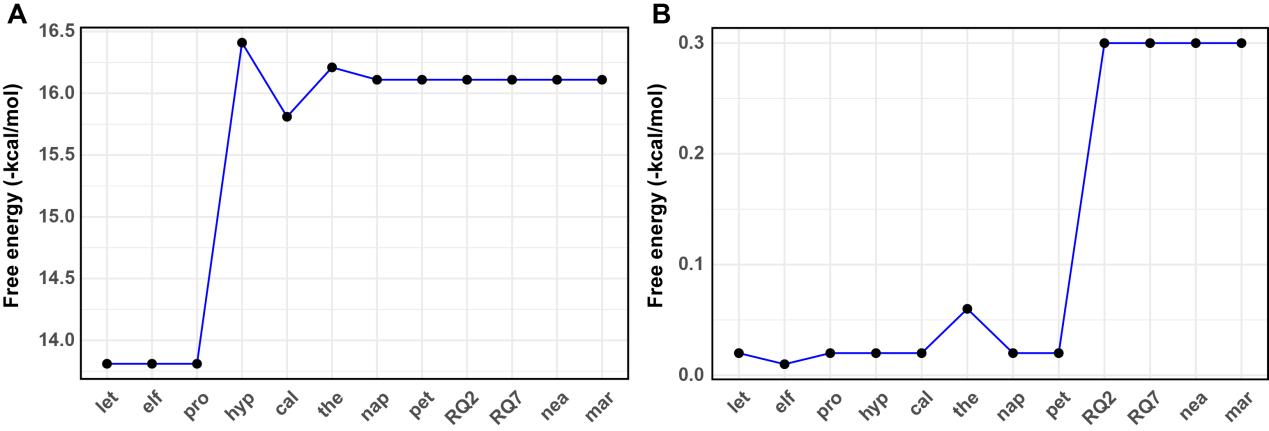

**Figure s3. Prediction of Free Energy for Two Stable G4s Identified in *Thermotoga*.** Prediction of free energy for two stable G4s identified in Thermotoga. (A) G4 structure at positions 1025-1054 nts. (B) G4 structure at positions 1474-1493 nts. Free energy was estimated using the RNAFold tool, with values influenced by nucleotide length and composition.
