## Supplementary material for "G-Quadruplex Structures in 16S rRNA Regions Correlate with Thermal Adaptation in Prokaryotes": Graphic abstract

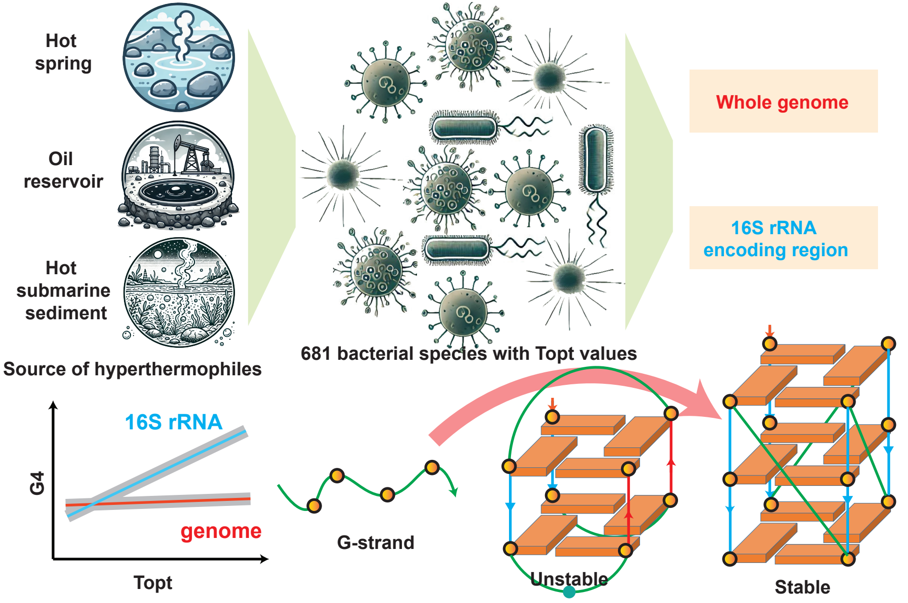


G4 positively correlates with Topt in 681 bacterial species, and the G4 structural stability and integrity in the 16S rRNA are evolutionarily enhanced in hyperthermophiles.
